# Multimodal mechanosensing enables treefrog embryos to escape egg-predators

**DOI:** 10.1101/2020.09.18.304295

**Authors:** Julie Jung, Shirley J. Serrano-Rojas, Karen M. Warkentin

## Abstract

Mechanosensory-cued hatching (MCH) is widespread, diverse, and improves survival in many animals. From flatworms and insects to frogs and turtles, embryos use mechanosensory cues and signals to inform hatching timing, yet mechanisms mediating mechanosensing *in ovo* are largely unknown. The arboreal embryos of red-eyed treefrogs, *Agalychnis callidryas,* hatch prematurely to escape predation, cued by physical disturbance in snake attacks. When otoconial organs in the developing vestibular system become functional, this response strengthens, but its earlier occurrence indicates another sensor must contribute. Post-hatching, tadpoles use lateral line neuromasts to detect water motion. We ablated neuromast function with gentamicin to assess their role in *A. callidryas*’ hatching response to disturbance. Prior to vestibular function, this nearly eliminated the hatching response to a complex simulated attack cue, egg-jiggling, revealing that neuromasts mediate early MCH. Vestibular function onset increased hatching, independent of neuromast function, indicating young embryos use multiple mechanosensory systems. MCH increased developmentally. All older embryos hatched in response to egg-jiggling, but neuromast function reduced response latency. In contrast, neuromast ablation had no effect on timing or level of hatching in motion-only vibration playbacks. It appears only a subset of egg-disturbance cues stimulate neuromasts; thus embryos in attacked clutches may receive uni- or multimodal stimuli. *A. callidryas* embryos have more neuromasts than described for any other species at hatching, suggesting precocious sensory development may facilitate MCH. Our findings provide insight into the behavioral roles of two mechanosensory systems *in ovo* and open possibilities for exploring sensory perception across taxa in early life stages.

**SUMMARY:** Red-eyed treefrog embryos use both their lateral line and vestibular systems to sense the disturbance cues in egg-predator attacks that inform escape-hatching decisions.

## INTRODUCTION

Animals ubiquitously use vibration to inform their behavior (Cocroft et al., 2014; Hill et al., 2019; Hill, 2009). Most research in this area focuses on vibrational communication signals in the contexts of mate choice, competition, and parental care (Cocroft et al., 2014). These signals are intentionally produced and generally operate between conspecifics. In addition, passively produced, unintentional vibrational cues can inform behavior. A smaller subset of research examines how animals use such incidental vibrations and other physical disturbance cues, generated either abiotically or biotically, to express context-appropriate behaviors. Conspecifics can generate disturbance cues (Doody et al., 2012), but other sources abound, such as weather (Marquez et al. 2016) and, more commonly, predators and prey (Bacher et al., 1997; Brownell and Leo van Hemmen, 2001; Castellanos and Barbosa, 2006; Oberst et al., 2017; Pfannenstiel et al., 1995; Roberts, 2018). Physical disturbances generated incidentally as animals move might be especially useful as cues because they can be difficult to conceal in a predator-prey context. Even embryos can sample incidental disturbance cues from their environments (Warkentin 2005; Warkentin, 2011a; Warkentin et al., In press). Like later life stages, embryos use physical disturbance cues from abiotic sources such as rainfall (Roberts, 2001), or biotic sources such as conspecifics (Endo et al., 2019; Noguera and Velando, 2019), hosts (Wang et al., 2012; Whittington and Kearn, 1988; Whittington and Kearn, 2011), and predators (Doody and Paull, 2013; Warkentin, 2005). From flatworms and insects to frogs and turtles, embryos of all sorts use disturbance cues to inform their hatching timing, yet the mechanisms mediating vibration and other mechanosensing *in ovo* are largely unknown.

The red-eyed treefrog, *Agalychnis callidryas*, is an excellent species in which to study how embryos detect and respond to physical disturbance or vibrations. As adults, these arboreal amphibians lay eggs on plants overhanging rainforest ponds. Usually within seven days, embryos hatch and fall into the water below, where they will develop as tadpoles, but they can hatch prematurely to escape from threats to eggs, including pathogens, flooding, and terrestrial egg predators (Warkentin and Caldwell, 2009). This escape-hatching response changes dramatically over development. The earliest observed hatching occurs at 3 days post-oviposition in response to strong hypoxia cues, but the onset of predator-induced, mechanosensory-cued hatching (MCH) does not occur until the following day (Warkentin et al., 2017). During the period of development between the onset of hatching responses to hypoxia and to mechanosensory cues, embryos are clearly able to hatch to escape threats. Nonetheless, they do not use this ability to flee from predators. Hatching ability is, therefore, not the sole constraint limiting the onset of escape-hatching responses to predator attacks.

The embryos of *Agalychnis callidryas* clearly use cues in multiple sensory modalities, including vibration (Warkentin 2005, Warkentin et al. 2019), hypoxia (Rogge and Warkentin, 2008) and light level (Güell and Warkentin, 2018), to inform hatching. In sensing physical disturbances, these embryos might use one or multiple mechanosensory systems, either to perceive different cue components available in attacks or as potentially redundant or synergistic sensors of the same cue component. As a first step toward determining the role of mechanosensory system development in the ontogeny of predator-induced hatching ability, we examined a general vertebrate motion sensor, the vestibular system of the inner ear (Jung et al., 2019). When otoconial organs in the developing vestibular system become functional, MCH increases substantially, but its earlier occurrence at a low level indicates that vestibular mechanoreceptors cannot be the only sensors that enable hatching when eggs are physically disturbed (Jung et al., 2019).

The mechanosensory lateral line system is found in all fishes and aquatic life stages of amphibians and serves to detect movement, vibrations, and pressure gradients in the surrounding water. It is comprised of mechanoreceptive neuromast sensory organs with hair cells that are sensitive to local water displacements (Lannoo, 1999) and are similar in morphology and function to hair cells in the auditory and vestibular system of vertebrates (Mogdans, 2019; Roberts et al., 1988). The lateral line system and vestibular system are responsive to many of the same stimulus fields (Braun and Coombs, 2000) and the hair cells of the lateral line and inner ear even have comparable thresholds of pressure detection (Van Netten, 2006). When something moves in the water, it creates water motion that deflects the ciliary bundles of the hair cells of the neuromasts, which opens mechanically gated ion channels (Harris et al., 1970; Sand et al., 1975). This stimulus information is sent to the brain, helping fishes and aquatic amphibians orient in currents (Elepfandt, 1982; Montgomery et al., 1997), maintain position within a school (Partridge and Pitcher, 1980), find prey (Bleckmann, 1980; Hoekstra and Janssen, 1985; Montgomery and Macdonald, 1987; Pohlmann et al., 2004), and detect predators (Montgomery, 1989; Schwalbe et al., 2012). Fishes and aquatic amphibians are able to detect low-level water motion with both the lateral line system and the inner ear, but the relative roles of these sensory systems is unclear (Karlsen and Sand, 1987).

Here we investigate the potential contribution of the embryonic lateral line system to sensing physical disturbance cues that inform escape-hatching decisions of *A. callidryas* embryos during snake attacks. When snakes bite, bump, and pull at eggs, embryos may receive complex mechanosensory stimuli, including motion, pressure, and tactile elements (Jung et al., 2019). The flow of perivitelline fluid, which constantly circulates around embryos, may also be altered as snakes stretch or squash egg capsules, thereby altering input to the lateral line system in the egg. However, not all embryos in an attacked clutch receive these complex cues. Embryos more distant from the snake may receive only whole-egg motion cues, transmitted through the clutch, but it is unclear if or how such whole-egg motion could stimulate the lateral line system (Jung et al., 2019). To assess the role of the lateral line and vestibular systems in sensing complex mechanosensory stimuli, we used a simulated attack cue (jiggling eggs in a standardized fashion with a blunt probe; Warkentin et al., 2017). We tested embryos with functional and inactivated lateral line systems, before and after the onset of vestibular function. To assess if egg motion alone stimulates the lateral line system, we used a custom-made vibration playback system to shake eggs in trays without providing concurrent tactile or egg deformation cues (Warkentin et al., In press; Warkentin et al., 2019) and tested embryos with functional and inactivated lateral line systems. Thus, we assessed the role of two mechanosensory systems, in two disturbance cue contexts, in a critical anti-predator behavior demonstrated by developing embryos.

## MATERIALS AND METHODS

### Egg clutch collection and care

We collected young (0–1 d old) *A. callidryas* egg clutches and the leaves on which they were laid from the Experimental Pond in Gamboa, Panama (9.120894 N, 79.704015 W) and brought them to a nearby laboratory (at ambient temperature and humidity) at the Smithsonian Tropical Research Institute. We mounted clutches on plastic cards for support, positioned them over aged tap water in plastic cups, then placed them inside plastic bins with screen lids connected to an automatic misting system (Mist King, Jungle Hobbies, www.mistking.com) set to mist clutches with rainwater at regular intervals to maintain hydration. Most clutches are laid between 10 pm and 2 am, so we assigned embryos to daily age-classes and reported developmental timing starting from midnight of their oviposition night (Warkentin, 2002; Warkentin, 2005). All tested individuals were staged based on external morphological markers (Warkentin, 2017) and were morphologically normal, in developmental synchrony with siblings in their clutch, and in intact, turgid eggs at the start of experiments. Hatchlings from gentamicin-treated eggs were reared in the laboratory under ambient conditions until hair cells regenerated (Hernández et al., 2007; Ma et al., 2008; Thomas et al., 2015; Williams and Holder, 2000), within 3 d of hatching. All hatchlings were returned to the pond from which they were collected. This research was conducted under permits from the Panamanian Environmental Ministry (SC/A-10-18 and SE/A-42-19) and approved by the Institutional Animal Care and Use Committee of the Smithsonian Tropical Research Institute (2017-0601-2020-2).

### Vestibulo-ocular reflex (VOR) measurement

To assess vestibular function, we measured roll-induced VOR of newly hatched tadpoles or manually decapsulated embryos using previously described methods (Horn and Sebastian, 1996; Horn et al., 1986; Jung et al., 2019). We placed hatchlings in a close-fitting tube of water and used a custom-made tadpole rotator to roll them about their body axis 180° in each direction, photographing them in frontal view each 15° (Jung et al., 2019). This method allowed us to measure vestibular function rapidly and without anesthesia. From each photograph, we measured right and left eye angle and body axis angle using ImageJ (Schneider et al., 2012). From each angular measurement series, we constructed an individual VOR curve using a sine-fitting function in Python (Version 2.7.9, Build 1, Python Software Foundation). The peak-to-peak amplitude of the curve fit is the measured VOR magnitude.

### Lateral line system visualization with 4-di-2-ASP

To visualize hair cells in neuromasts, newly hatched or decapsulated tadpoles were immersed in a 500 μM solution of the fluorophore 4-(4-diethylaminostyryl)-1-methylpyridinium iodide (4-di-2-ASP, Sigma D-3418, sigmaaldrich.com) for five minutes at room temperature in the dark. Tadpoles were then rinsed in aged tap water to remove excess fluorophore and placed in a shallow petri dish or snout-up in a modified pipette mount. This method allowed us to label neuromast hair cells and photograph them in live embryos without anesthesia or treatment with a paralytic. We used an Olympus stereoscope equipped with epifluorescence (GFP filter set) to photograph each individual. We considered neuromasts showing fluorescence to be functional and a complete lack of fluorescence to indicate neuromast inactivation. Fluorescence gradually faded within hours and disappeared within a day of treatment. Treated individuals developed normally compared to untreated controls.

### Lateral line system knockout with gentamicin

To test the role of the lateral line system in predator-induced, mechanosensory-cued escape-hatching behavior, we used the ototoxic aminoglycoside antibiotic gentamicin to ablate neuromast function (Kroese and van den Bercken, 1982; Montgomery et al., 1997; Van Trump et al., 2010). At embryonic age 3 d, we removed individual eggs from clutches and placed groups of eight siblings into hexagonal polystyrene weigh boats (4.5 cm across x 1 cm depth). We added 1 mL of rainwater to each boat, partially submerging eggs, and waited 1 h for water absorption to increase and standardize turgidity before treatment. To begin treatment, we removed the remaining water and added a fresh 1mL of rainwater into each boat. Control eggs remained in rainwater, while treatment eggs received a timed dosage series of gentamicin sulphate solution-10 (MP Biomedicals**™**, fishersci.com) mixed into the rainwater to gradually increase the concentration (starting dose 2 μL, + 2 μL at 3 h, + 2 μL at 6 h, + 0.5 μL at 9 h, + 0.5 μL at 10.5 h). The final concentration was the minimum dosage that effectively blocked lateral line system function, determined based on pilot experiments. The gradual increase in concentration was necessary to maintain egg turgor and membrane integrity, as needed for hatching assays, also based on pilot experiments. At 13 h, we extracted 0.5 mL of solution from each boat to increase air exposure of eggs and ensure sufficient oxygen availability for these terrestrial eggs as they continued developing in the boats until hatching-response tests were carried out. Only eggs that maintained normal turgidity until testing were used to assess embryo responses to mechanosensory cues. We confirmed lateral line system blocking with 4-di-2-ASP staining in each experiment (Fig. 1A-B). To assess if gentamicin damaged hair cells in embryonic ears, as suggested previously (Bagger-Sjoback, 1997; Simmons et al., 2004; Van Trump et al., 2010; Yan et al., 1991), we compared VOR of stage-matched (within one developmental stage of each other) control and gentamicin-treated siblings, across all experiments described below (Fig. 1C).

**Figure 1.**
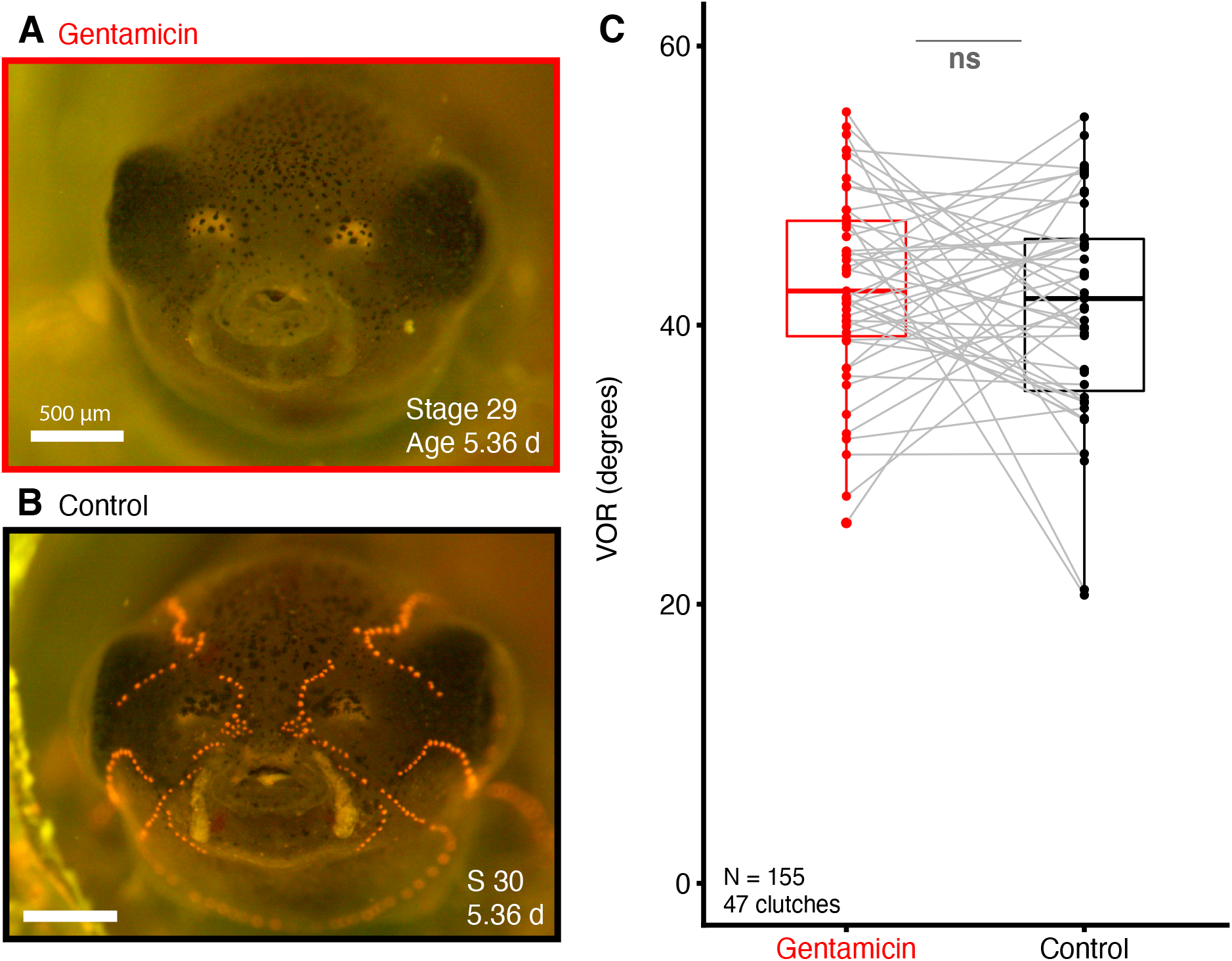
Gentamicin ablates lateral line system function with no effect on vestibular system. (A) Representative gentamicin-treated individual, showing complete ablation of functional neuromasts, and (B) age- and stage-matched control sibling, showing functional neuromasts (orange). (C) VOR amplitude of gentamicin-treated and control *A. callidryas*. Gray lines connect points representing stage-matched siblings, in the same or adjacent developmental stages, across treatments. ns P > 0.05.

### Hatching-response test: manual egg jiggling

To assess the MCH response of gentamicin-treated (Fig. 1A) and control (Fig. 1B) embryos across ontogeny, we performed a standard egg-jiggling assay (simulated attack), using a blunt probe to manually jiggle eggs in an intermittent pattern for 5 min (Jung et al., 2019; Warkentin et al., 2017). This simulated attack includes complex motion and tactile elements, and transient deformation of egg membranes. We tested 1202 individual embryos from 60 clutches (at least 5 individuals per treatment per clutch) from July 2 to August 5, 2019. We moved sibling pairs of eggs (treatment and control individuals) into their own weigh boats with a drop of water and manually jiggled them with a blunt metal probe, alternating 15 s of stimulation and 15 s of rest (i.e., stimulating eggs in a pair alternately) for 5 minutes or until the embryo hatched.

We exposed embryos to jiggling cues during three developmental periods (Fig. 2A): (1) across the onset of vestibular function (N=544 individuals later subdivided based on VOR measurements; range, mean ± SEM age: 4.03–4.36 d, 4.19 ± 0.003 d; stage 27–28, 27.19 ± 0.017), (2) after vestibular function was well-established (N=360 individuals; 4.54–5.03 d, 4.74 ± 0.007 d; stage: 27–30, 28.99 ± 0.010), and (3) closer to spontaneous hatching (N=298 individuals, age: 5.29–5.56 d, 5.39 ± 0.003 d, stage: 29–32, 30.44 ± 0.046). For the youngest age group, we measured the VOR of all individuals that hatched (N=67 individuals) and checked for lateral line system blocking of all hatched, gentamicin-treated individuals (N=3). We used VOR measurements to determine vestibular function onset times, dividing siblings into subsets tested before and after VOR onset. For the two older groups, we measured VOR in at least one gentamicin-treated and one control individual per clutch (N=111 individuals). We used 4-di-2-ASP staining to check for lateral line system blocking in at least one individual per gentamicin-treatment boat with hatching and visualized neuromasts in one untreated control per testing session (i.e., the period of hours over which a group of trays were tested; N=31 individuals).

**Figure 2.**
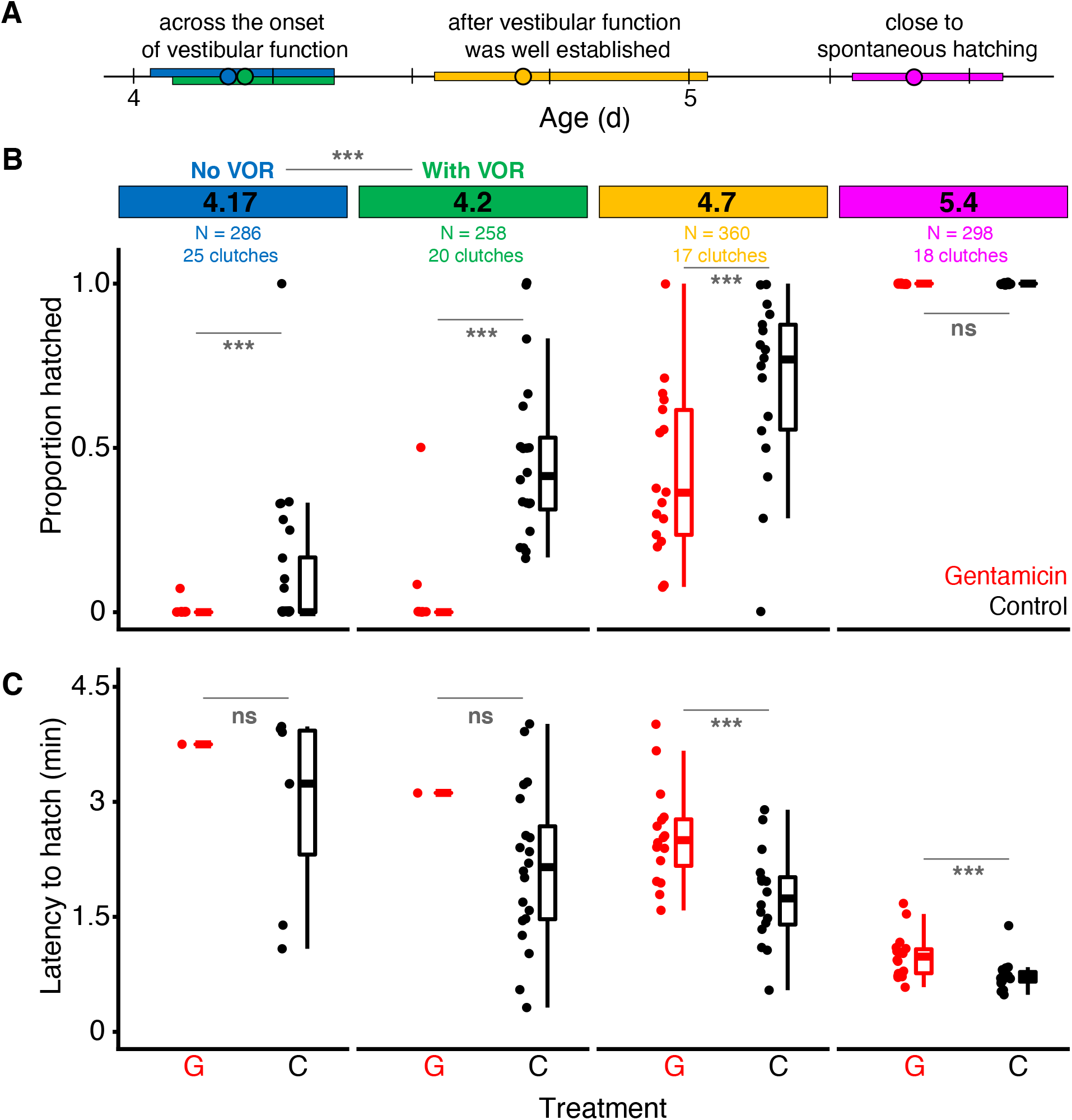
Ontogeny of the hatching response to egg-jiggling by gentamicin-treated and control *A. callidryas* embryos, across and after the onset of vestibular function (indicated by VOR). (A) Developmental timeline of testing periods, with mean ages (points) and ranges (rectangles) color coded, (B) hatching responses, and (C) latencies. Data points are clutch-means and boxplots show means, quantiles, and 1.5*IQR across clutches, within treatment × age category; ns P > 0.05, *** P < 0.001.

For the youngest developmental period, we determined the start of testing for each sibship based on external developmental markers (Warkentin, 2017). We began testing early in stage 27, near the mean first appearance of MCH across clutches (Warkentin et al., 2017) and shortly before the onset of vestibular function (Jung et al., 2019), then stimulated pairs of siblings sequentially until VOR was visible in tested hatchlings. After analysis of VOR tests, we estimated the onset of vestibular function for each clutch as the first time point at which any individual from the clutch showed a VOR > 10°. Applying this criterion generated two groups: “no VOR” (N=286 individuals; age 4.03–4.36, 4.17 ± 0.005 d; all stage 27) and “with VOR” (N=258 individuals, age: 4.07–4.36, 4.20 ± 0.005; stage 27–28, 27.23 ± 0.027). These two groups differ significantly in age (Wilcoxon Rank Sum: z=3.8, P=0.00007). In prior work using manual egg-jiggling at stages 26–29 and measuring the VOR of every individual, the hatching response of individuals “without vestibular function” was similarly low (23%) using criteria of <1° or <10°, and it increased substantially as VOR increased to ca. 30–40° (Jung et al., 2019). Here, using the VOR of measured individuals to estimate the presence/absence of vestibular function in their unmeasured siblings allowed us to test for MCH in many more individuals than we could measure for VOR, but entails some classification errors. Applying our criterion to a prior dataset (Jung et al., 2019) we expect an estimated level of 14% false positives and 22% false negatives, reducing the chance of detecting an effect of vestibular function on hatching in the current data.

For each individual, we recorded hatching response (Fig. 2B) and latency (seconds from stimulus onset) or failure to hatch after 5 min of post-stimulus observation (Fig. 2C). Proportion hatched was calculated per treatment per clutch.

### Hatching-response test: vibration playbacks

To provide a motion-only stimulus, without concurrent tactile cues, we performed vibration playbacks to groups of embryos held in custom-made egg-trays (Warkentin et al., In press; Warkentin et al., 2019). Trays held up to 15 eggs (>8, 11.79 ± 0.25 eggs per tray) in individual funnel-shaped spaces, allowing hatched tadpoles to slide through the tray to water below. We tested 1132 individuals in 96 trays from 63 clutches from July 25 to August 7, 2019 (age: 5.40– 5.75, 5.56 ± 0.009 d, stage: 30–33.67, 31.27 ± 0.080). We aimed to test embryos of the same age as the 5 d individuals tested with jiggling cues, but due to logistic constraints playback embryos were, on average, 4 hours older than jiggled embryos. Embryos were treated with gentamicin or as rainwater controls at age 3 d, as described above, and moved to individual spaces in egg-trays early at embryonic age 4 d, while eggs could be easily handled without inducing hatching. Full trays were maintained on racks over aged tap water in humidors until testing, with regular misting until age 5 d (Warkentin et al., In press; Warkentin et al., 2019).

We designed a synthetic low-frequency vibration stimulus to elicit high hatching rates, based on prior playbacks to 5 d embryos (Caldwell et al., 2009; Jung et al., 2019; Warkentin et al., In press; Warkentin et al., 2006b; Warkentin et al., 2017; Warkentin et al., 2019). We generated white noise in MATLAB and filtered it using a custom script (available upon request) to compensate for nonlinearities in the shaker transfer function and generate a frequency distribution similar to snake-attack vibrations (Caldwell et al., 2009), with high energy below 50 Hz and intensity dropping off above that (Fig. 3A). The temporal pulse pattern included a 3-pulse ‘primer’ and a series of seven 10-pulse groups separated by 30 s intervals of silence (Fig. 3B). Within each pulse group, the base temporal pattern had a cycle length of 2 s, consisting of 0.5 s pulses of vibration with roughly rectangular amplitude envelopes separated by 1.5 s intervals of silence (Fig. 3C). To record playback vibrations, we attached a small (0.14 g) AP19 accelerometer (AP Technology International B.V., Oosterhout, The Netherlands) to an egg-tray using dental wax (Warkentin et al., In press). Accelerometer output was routed through an AP5000/10 charge-to-voltage converter to a B&K 1704 signal conditioner (Brüel & Kjær, Nærum, Denmark), digitized with a Focusrite Scarlett 2i2 external sound card (focusriteplc.com), and recorded using Raven Pro 1.3 bioacoustics software (Cornell University Laboratory of Ornithology, Ithaca, NY, USA) on a Macbook Air computer. The RMS amplitude of the playback vibrations, excluding intervals of silence, was 6.5 ms^−2^.

**Figure 3.**
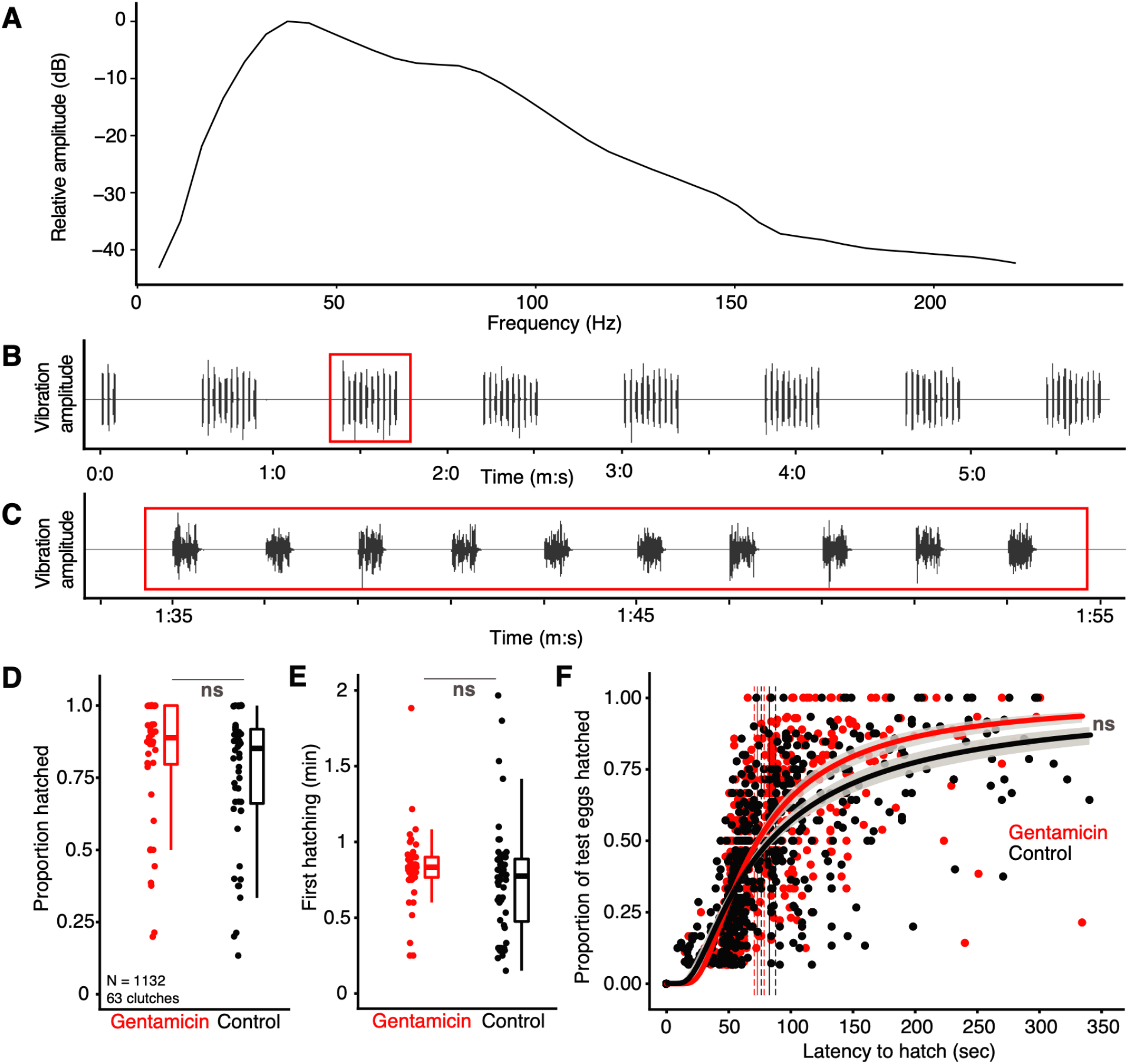
Hatching responses of gentamicin-treated and control *A. callidryas* embryos (mean age 5.56 d, stage 31.3) to motion-only vibration playback. (A) Frequency spectrum from recording of playback stimulus, (B) waveform of entire stimulus, with (C) waveform of a single pulse group in red box, showing the base temporal pattern (0.5 s duration, 1.5 s interval). (D) Proportion of embryos hatched and (E) latency for the first individual to hatch, after stimulus began. Points are values per tray, boxplots show means, quantiles, and 1.5*IQR across trays. (F) All individual hatching latencies (points) in playbacks, with Weibull curve fits and 95% CI (shading). Vertical solid lines indicate the mean latency by which 50% of individuals within treatments had hatched (estimated from bootstrap), dashed lines indicate the 95% CI. ns P > 0.01.

Playback methods followed published work detailing vibration presentation via egg-trays (Warkentin et al., In press; Warkentin et al., 2019). For testing, we clamped egg-trays holding embryos to a custom-made interface on a rigid post attached to an electrodynamic minishaker (Model 4810; Brüel & Kjær, Nærum, Denmark). The shaker, post, and tray were horizontally leveled, with foam supports under the post and tray edge and water under the embryos. Thus, embryos were moved horizontally and hatchlings fell into the water below. Shaker output was controlled by Audacity 2.1.0 on a 2014 Macbook Air, via a custom-made amplifier (E. Hazen, Boston University Electronic Design Facility). We recorded any hatching induced by the set-up procedure (N=3 individuals), then allowed five minutes for acclimation before starting the playback (N=38 individuals hatched during acclimation). Individuals that hatched before the stimulus started were not considered part of the test. We noted if and when (to the nearest second) each embryo hatched during stimulus playback and 3 minutes of post-playback observation (Fig. 3D-F). We then immediately (within 20 min) measured VOR of a subset of individuals, including a pair of gentamicin-treated and control individuals from 32 clutches (plus 14 clutches with unpaired data). We confirmed hatching competence of embryos remaining unhatched by manually stimulating them with a blunt metal probe (N=251 individuals). We also used 4-di-2-ASP staining to confirm lateral line blocking in at least one individual per treated tray with hatching and to visualize neuromasts in one control per testing session (N=43 treated, 5 untreated individuals). We staged 3 haphazardly selected hatchlings from each tray (Warkentin, 2017).

### Lateral line system ontogeny

To determine at what point during lateral line development the hatching response to jiggling cues begins, we compared the ontogeny of the lateral line system and of MCH, using a developmental series of 79 embryos from 25 clutches from August 6–11, 2018. For each individual, we first tested for MCH, using the same manual egg-jiggling stimulus as in the first hatching response test (Warkentin et al., 2017). Test clutches were transported just prior to hatching competence and tested in an air-conditioned laboratory of the Smithsonian Tropical Research Institute in Gamboa. We recorded latency to hatch and embryos that remained unhatched after 5 min were manually decapsulated using sharp forceps. We staged embryos under a stereoscope (Warkentin, 2017). Neuromasts were then stained using 4-di-2-ASP as described above and photographed in in frontal and dorsal views. The frontal view showed neuromasts in the ventral, oral, infraorbital, nasal, and supraorbital lines (Fig. 4A), while the dorsal view showed neuromasts in the nasal, supraorbital, middle, and dorsal lines (Fig. 4B). All neuromasts within each line were labeled and counted using ImageJ software (Schneider et al., 2012). We took several photos in each view, in different focal planes, framing, and magnification, then counted neuromasts in each line from the photo(s) that showed them best. Counts from supraorbital and middle neuromast lines were extrapolated from one side, assuming bilateral symmetry within individuals. When lines extended across multiple photos, we used landmarks to avoid double-counting or missing neuromasts. We averaged counts per line from two independent counters and summed them across lines to estimate the total number of neuromasts per individual (Fig. 4C, Table S1). We also measured tadpole total length from dorsal images (Fig. 4D).

**Figure 4.**
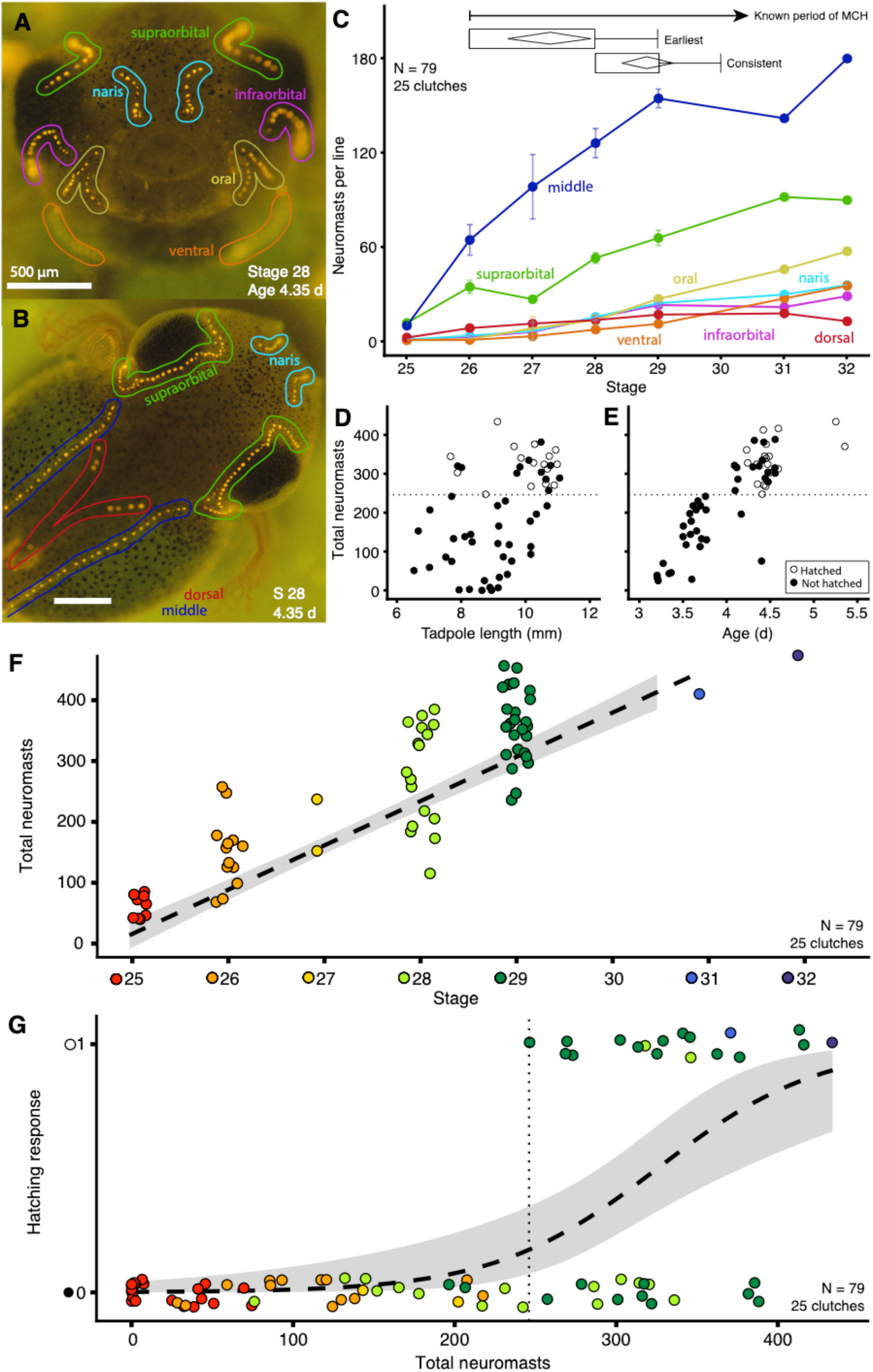
Lateral line system ontogeny and development of the hatching response of *A. callidryas* embryos to egg-jiggling. Representative (A) frontal and (B) dorsal views of hatchlings stained with 4-di-2-ASP to show neuromasts, with lines color-labeled. (C) Mean number of neuromasts in each line (bilaterally) across development, ± SEM. Inset shows previously reported (Jung et al., 2019; Warkentin et al., 2017) stages when MCH is first and consistently expressed; diamonds represent means and 95% confidence intervals, box and whiskers show interquartile range (IQR) and extent of data to ± 1.5×IQR. Total number of neuromasts increased with (D) total length (χ^2^=5.2, df=1, P=0.02319), (E) age (χ^2^=62.4, df=1, P<2.9e-15), and (F) developmental stage (χ^2^=6263.8, df=1, P<2.2e-16; linear regression and 95% CI indicated). (G) Hatching response increased with neuromast number; predicted curve fit (dashed line) and 95% CI (shading) from binomial generalized linear mixed model are indicated. For (G,H) categorical variables are jittered to show data points and stages are color-coded. Dotted lines (D,E,G) show the threshold number of neuromasts at which hatching began.

### Statistics

All statistical tests were carried out in the R statistical environment (version 3.6.2, R Development Core Team 2019, http://www.r-project.org) in RStudio (version 1.2.5033, RStudio Team 2019). We used generalized linear mixed models within the ‘lme4’ package (Bates et al., 2015) with clutch as a random effect and likelihood ratio tests to compare nested models for fixed effects and interaction effects on the number of neuromasts (error distribution: negative binomial), hatching responses (binomial), and first hatching latency (gamma). We used the fitconds function within the ‘fitplc’ package to plot curve fits for all hatching latencies (weibull) within trays (Duursma and Choat, 2017).

## RESULTS

### Lateral line system knockout with gentamicin

Our gentamicin treatment completely inactivated all neuromasts with no evidence of differences in the VOR of stage-matched gentamicin treated and non-treated siblings (χ^2^_1_=2.3, P=0.1302, Fig. 1).

### Hatching-response test: egg jiggling

Across the onset of VOR (mean age 4.17 vs. 4.2 d), both lateral line and vestibular function strongly and independently increased the likelihood of hatching in response to egg-jiggling (gentamicin: χ χ^2^_1_=88.2, P<2.2e-16; VOR: χ^2^_1_=30.4, P=3.5e-8; interaction: χ^2^_1_=0.7, P=0.4, Fig. 2A-B). In embryos lacking vestibular function (mean age 4.17 d), gentamicin reduced the hatching response significantly from 10% to 1% (χ^2^_1_=14.4, P=0.0001, Fig. 2B). Just after the onset of VOR, gentamicin still reduced hatching significantly (41% vs. 2%; χ^2^_1_=73.8, P<2.2e-16, Fig. 2B). While both age and neuromast function increased hatching, the gentamicin effect on MCH decreased with age (age: χ^2^_3_=200.1, P<2.2e-16; gentamicin: χ^2^_1_=105.7, P<2.2e-16; interaction: χ^2^_3_=19.9, P=0.0002; Fig. 2B). At 4.7 d, gentamicin still significantly reduced hatching (70% vs. 41%; χ^2^_1_=35.4, P=2.7e-09) but at 5.4 d, close to spontaneous hatching, all jiggled eggs hatched, with or without functional neuromasts (Fig. 2B).

Latency to hatch decreased with age and lateral line function (age: χ^2^_3_=79.2, P<2.2e-16; gentamicin: χ^2^_1_=36.9, P=1.266e-09; interaction: χ^2^_3_=3.8, P=0.29; Fig. 2C). The low hatching response of gentamicin-treated embryos before and just after the onset of VOR limited our sample of latency, reducing statistical power for comparisons (Fig. 2C). However, gentamicin treatment increased latency to hatch at both 4.7 d (χ^2^_1_=17.1, P=3.5e-05) and 5.4 d (χ^2^_1_=21.2, P=4.1e-06) with no indication that the effect decreased with age (interaction: χ^2^_1_=2.84, P=0.09; Fig. 2C).

### Hatching-response test: vibration playbacks

To assess if egg motion alone can stimulate the lateral line system, we used vibration playbacks to shake gentamicin-treated and untreated embryos held in custom-made egg trays. In these playbacks, lateral line function had no effect on the hatching response of embryos (χ^2^_1_=1.8, P=0.1787, Fig. 2D) or timing of hatching (χ^2^_1_=1.1, P=0.2986, Fig. 2E-F).

### Lateral line system ontogeny

The number of neuromasts in all seven lines increased with developmental stage (Fig. 4C, Table S1). Total number of neuromasts increased with size (Fig. 4D), age (Fig. 4E) and, most strongly, developmental stage (Fig. 4F). From stage 27 to 29, the number of neuromasts more than doubled (Fig. 4F). Both developmental stage (χ^2^_1_=50.9, P=9.5e-13) and the total number of neuromasts (χ^2^_1_=15.7, P=7.5e-5) were significant predictors of hatching in egg-jiggling experiments (Fig. 4G). When embryos had fewer than a threshold number of neuromasts (247), none hatched, whereas embryos with more neuromasts often hatched.

## DISCUSSION

We demonstrate that the developing lateral line and vestibular systems both contribute to escape-hatching of red-eyed treefrog embryos, and that the roles of these sensors change during development and vary with disturbance cue type.

### Lateral line system knockout with gentamicin

Using gentamicin, we achieve complete lateral line ablation in *A. callidryas* embryos with no evidence of vestibular system damage (Fig. 1). Others found evidence that gentamicin can cause vestibular system damage when administered intramuscularly (Bagger-Sjoback, 1997; Yan et al., 1991) or by direct immersion post-hatching (Simmons et al., 2004; Song et al., 1995; Van Trump et al., 2010). Our study is the first to investigate the effects of gentamicin administered incrementally to embryos developing *in ovo*, which was necessary to avoid water loss from eggs and, later, test for hatching responses. The gradual passage of gentamicin across the vitelline membrane into the perivitelline space while eggs were in the treatment bath, and the potential loss of gentamicin from older eggs maintained in egg-trays before vibration playbacks, means we do not know the precise concentrations embryos were exposed to over time. However, our longest exposure durations and peak exposure concentrations exceeded those in previous studies (Simmons et al., 2004; Song et al., 1995; Van Trump et al., 2010), supporting that for some animals under some exposure conditions gentamicin can selectively damage hair cells in the lateral line without impairing the function of hair cells in the vestibular system. Lateral line neuromasts are directly exposed to the fluid bathing an embryo, but hair cells of the inner ear are not, and the barriers protecting these internal cells likely change with development.

Notably, we found no evidence for ear damage in early stages, at the onset of vestibular function, and also in more developed 5 day old embryos. Immature and recently regenerated hair cells are resistant to aminoglycoside antibiotics (Dai et al., 2006; Hashino and Salvi, 1996; Murakami et al., 2003; Van Trump et al., 2010). Thus, in the youngest tested embryos, vestibular hair cells may have had only brief gentamicin exposure after maturing to a point of vulnerability. In the lateral line system, neuromasts began to regain fluorescence, when stained with DiAsp, within hours of hatching and cessation of gentamicin exposure. We suspect this was due to maturation of gentamicin-resistant developing hair cells. Embryos tested at 5 days would have had functional hair cells in their ears for over 24 h (29.8 h from mean onset of VOR to mean testing age) under gentamicin treatment, yet we also found no evidence for vestibular damage in these older embryos. This suggests that, compared to more mature tadpoles (Bagger-Sjoback, 1997; Simmons et al., 2004; Van Trump et al., 2010; Yan et al., 1991), embryonic anatomy may better protect otic hair cells against externally administered gentamicin.

### Hatching-response test: manual egg jiggling

In embryos lacking vestibular function, a significant effect of gentamicin treatment on hatching response reveals that the onset of lateral line function precedes the onset of vestibular function and plays a key role in very early MCH (Fig. 2B). Immediately following the onset of VOR, the lateral line system continues to contribute strongly to risk assessment and hatching. The higher hatching rate in VOR-positive animals (Fig. 2B) is consistent with a key role of otoconial organs in embryonic vibration sensing (Jung et al., 2019). However, at the onset of MCH neither lateral line nor vestibular function alone enabled a strong response; the multimodal combination of input from both senses greatly increased the likelihood of hatching in a simulated attack (Fig. 2B).

Soon after *A. callidryas* gain hatching competence, the lateral line system plays a critical role in sensing and responding to predator cues. We tested the effect of gentamicin on jiggling-induced hatching in two later periods to determine if the dependence of MCH on lateral line function changes developmentally. In *A. callidryas*, embryos hatch spontaneously from 5–7 d, while younger embryos almost never hatch if undisturbed (Güell and Warkentin, 2018; Hite et al., 2018; Warkentin, 2000; Warkentin et al., 2001). From age 4 to 5 d, embryos become more likely to escape during snake and wasp attacks (Gomez-Mestre and Warkentin, 2007; Warkentin, 1995; Warkentin, 2000) and to hatch in vibration playbacks (Jung et al., 2019; Warkentin et al., In press; Warkentin et al., 2019). Over this same developmental period, our results show that MCH responses become less dependent on multimodal input from the lateral line plus vestibular system. This might reflect increasing strength of vestibular input as embryo ears develop (Jung et al., 2018).

When animals use multimodal sensory integration to inform behavior, a sensor may contribute to a response without being required for its occurrence (Angelaki and Cullen, 2008). We examined latency from stimulus onset to hatching response (Fig. 2C) as a potentially more sensitive indicator of lateral line system contributions to embryo decisions, since latency affects escape success during predator attacks (Almanzar and Warkentin, 2018; Chaiyasarikul and Warkentin, 2017). We found that even embryos near the stage of spontaneous hatching, when they have a very strong vestibular-mediated hatching response, still use lateral line input to accelerate their response to simulated attack cues (Fig. 2C).

### Hatching-response test: vibration playbacks

The fact that lateral line function had no effect on the proportion of embryos hatching (Fig. 3D) or their latency to hatch (Fig. 3E) in response to motion-only vibration playbacks suggests that only a subset of more complex physical disturbances stimulate the lateral line system, while potentially any egg motion may stimulate the vestibular system. Thus, some embryos in attacked egg clutches receive unimodal vestibular stimulation while others receive multimodal vestibular and lateral line system input. Some may also receive input from cutaneous touch receptors (Blaxter, 1987; O’Brien et al., 2012). Given the role of lateral line system input in reducing hatching latency, as well as increasing hatching likelihood in younger embryos, this variation in the type(s) of sensory input an embryo receives under different predation contexts likely contributes to the variation in the likelihood and timing of hatching during attacks (Almanzar and Warkentin, 2018; Warkentin, 2000; Warkentin et al., 2006a). The variation in received mechanosensory stimuli within a clutch may also be a mechanism contributing to threat-sensitive embryo behavior (Ferrari and Chivers, 2009; Hughey et al., 2015; Mathis et al., 2008; Van Buskirk, 2016); since predators must touch eggs to eat them, the risk of mortality during attacks is likely higher for eggs receiving multimodal cues than for those receiving motion cues alone.

### Lateral line system ontogeny

In a recent study, we found hatching in response to egg-jiggling to begin, on average, at stage 27.3 and considered hatching “consistent” at stage 28.8 (Fig. 4C), the second time both of two tested siblings hatched (Jung et al., 2019; Warkentin et al., 2017). This known onset of MCH aligns ontogenetically with a rapid increase of neuromast number in developing embryos (Fig. 4C). The pattern of a neuromast-number threshold for hatching (Fig. 4G) suggests that lateral line system development may limit the onset of disturbance cue sensing and associated hatching behavior.

### Lateral line morphology in comparative context

The number of neuromasts at hatching and their proliferation after hatching varies greatly across taxa. Some fishes hatch with only two functional neuromasts, and early larvae simply float in the water column [e.g. flounder (Kawamura and Ishida, 1985), tuna (Kawamura et al., 2003), grouper (Mukai et al., 2006), catfish (Mukai et al., 2010)]. Subsequent rapid lateral line system development is correlated with behavioral changes (avoiding obstacles, feeding, migrating, settling, and surviving seasonal floods) and parallels rapid development of other sense organs such as eyes, ears, taste buds, olfactory epithelium (Kawamura and Ishida, 1985; Kawamura et al., 2003; Mukai et al., 2006; Mukai et al., 2010). Neuromast proliferation can also occur slowly. For instance, cod larvae hatch with five lateral body neuromasts, and only add one more by feeding onset, 2–3 weeks later (Blaxter, 1984). The developmental stage at which neuromast function begins differs among species, and appears related to hatchlings’ habitat and habits (Otsuka and Nagai, 1997). For instance, ayu hatch with 20 pairs of well-developed neuromasts and migrate downstream immediately upon hatching (Kawamura et al., 1983). In contrast, pale chub hatch without a single neuromast and remain in the spawning bed for 4 d, during which neuromasts develop rapidly; at emergence into the river, larvae are responsive to water flow and have nearly caught up to ayu in neuromast number (Kawamura et al., 1983). Across species, developmental increases in the number of neuromasts are closely linked to the ontogeny of mechanosensory-guided behavior (Blaxter and Fuiman, 1989; Kawamura and Ishida, 1985; Llanos-Rivera et al., 2014). However, neuromast size, shape, and hair cell polarity can also affect mechanosensory sensory function (Becker et al., 2016; Webb and Shirey, 2003).

Anatomically and physiologically, amphibian neuromasts resemble superficial neuromasts of teleost fishes (Metcalfe, 1985; Simmons et al., 2004). However, we know relatively little about their early development, except in a few species. At hatching, the salamander *Ambystoma mexicanum* has all 60 neuromasts, while in the frog *Lithobates (Rana) pipiens* most neuromasts have not yet formed; however, by the onset of feeding, the lateral line appears fully formed and functional in both species (Smith et al., 1988). Among amphibians, early lateral line system development has been characterized in detail in *Xenopus laevis* (Roberts et al., 2009; Shelton, 1970; Simmons et al., 2004; Winklbauer, 1989), but studies rarely distinguish hatching timing. Reported hatching stage varies such that *X. laevis* have between 0 (Carroll and Hedrick, 1974) and 14 neuromasts at hatching, but embryos with just 6–8 neuromasts swim into water currents and responsiveness increases over the next 10 h with an increase in neuromast number (Roberts et al., 2009). In our study system, the hatching response of *A. callidryas* embryos to disturbance begins only when they have hundreds of neuromasts (Fig. 4G). Despite the large increase in neuromast number, there appears to be little change in neuromast size (personal observation from confocal images by María José Salazar Nicholls and Julie Jung) or shape (Cohen et al., 2019) from 3 to 5 days post oviposition. It is not yet clear how physical disturbance of eggs stimulates the neuromasts of embryos inside them; however, induced perturbations of perivitelline fluid motion within the egg capsule may provide a weaker or less clear stimulus to the lateral line system than currents in open water do after hatching. Additionally, the cost of unnecessarily swimming into a current may be lower than that of hatching prematurely.

The number of neuromasts *A. callidryas* have at hatching is high, even compared to other anurans in late larval development (Lannoo, 1987). *A. callidryas* that hatched in response to egg-jiggling, at 7.7–11.0 mm total length, had neuromast counts of 323.6 ± 11.9 (range 247–434). For comparison, in a study of 36 other anuran species examined in late larval development, at their expected peak of neuromast numbers (Lannoo, 1987; Winklbauer, 1989), the most neuromasts reported was 332 in *Rana aurora*, at 25–27.5 mm snout-vent length (Lannoo, 1987). Since lateral line development depends on the size of the animal, and tadpoles tend to add neuromasts during post-hatching development (Fabrezi et al., 2012; Winklbauer, 1989), these other species presumably have fewer neuromasts at hatching. This suggests that *A. callidryas* have precocious lateral line development, as well as high neuromast numbers. However, we know nothing about lateral line ontogeny in other anurans with documented or suspected mechanosensory-cued escape-hatching behavior (Brown and Iskandar, 2000; Brown et al., 2010; Buckley et al., 2005; Chivers et al., 2001; Gomez-Mestre et al., 2008; Poo and Bickford, 2014; Sih and Moore, 1993; Smith and Fortune, 2009; Touchon et al., 2011). Egg predation might be a selective force favoring earlier or greater development of mechanosensory systems. A comparative analysis of lateral line and vestibular system development in relation to the distribution of MCH in anurans (Warkentin, 2011b; Warkentin, 2011a) would be informative. Assessing links between sensory morphology and embryo behavior could reveal patterns of functional variation and adaptive evolution and provide further insight into how embryos use sensory information.

Embryos across diverse taxa respond to physical disturbance cues from many ecologically relevant sources (Warkentin, 2011b; Warkentin, 2011a; Warkentin et al., In press). In contexts such as antipredator defense (Doody and Paull, 2013; Warkentin, 2005) and sibling hatching synchronization (Endo et al., 2019; Noguera and Velando, 2019), embryos’ ability to sense these cues is often essential for their survival, yet we know very little about the sensory systems that mediate MCH or even, in many cases, which elements of physical disturbance cues are most relevant. Herein, we show that to detect egg-disturbance cues *A. callidryas* embryos use their vestibular system, a sensory mechanism likely to be relevant across vertebrates, and their lateral line system, a mechanism which may contribute to MCH in other frogs and fishes. Our study serves as proof-of-concept for the feasibility of combining neuromast ablation *in ovo* with hatching-response tests, and it highlights the value of latency as a sensitive response variable in the context of multimodal mechanosensing. The methods we present open opportunities for comparative research into embryo sensory ecology with other species that hatch in response to mechanosensory cues. Our findings provide insight into the functional roles of two different mechanosensory systems prior to hatching and reveal new possibilities for exploring embryonic sensory perception across taxa.

## Acknowledgments

This research was funded by the National Science Foundation (IOS-1354072 to KMW and James Gregory McDaniel), and conducted under permits SC/A-10-18 and SE/A-42-19 from the Panamanian Ministerio de Ambiente and STRI IACUC protocol 2017-0601-2020-2. We thank J. Gregory McDaniel for collaboration in biotremology, including development of the tadpole rotator and egg-tray playback system, Jacqueline Webb and Andrea Simmons for advice on neuromast blocking, Estefany Caroline Guevara Molina, Luis Alberto Rueda Solano, and Henry Macías for help with egg jiggling and care, Casey Lam for assistance measuring eye angles and body angles from photographs, members of the Gamboa Frog Group at STRI and BU Egg Science Research Group for discussions of this research, and Jacqueline Webb and Pete Buston for helpful comments on the manuscript.

## Author Contributions

Conceptualization: J.J. and K.M.W.

Methodology: J.J., S.J.S.R., and K.M.W.

Software: J.J.

Validation: J.J., S.J.S.R., and K.M.W.

Formal analysis: J.J.

Investigation: J.J. and S.J.S.R.

Resources: K.M.W.

Data curation: J.J.

Writing – original draft: J.J.

Writing – review and editing: J.J., S.J.S.R., and K.M.W.

Visualization: J.J.

Supervision: K.M.W.

Project administration: K.M.W.

Funding acquisition: K.M.W.

---

**Table S1.** Lateral line system ontogeny. Mean (± SE) number of neuromasts in each line (bilaterally) of the lateral line system in *A. callidryas* embryos across the onset of mechanosensory-cued hatching.

| *Stage* | *n* | *Infraorbital* | *Supraorbital\** | *Naris* | *Oral* | *Ventral* | *Middle\** | *Dorsal* | *Bilateral Total* |
| --- | --- | --- | --- | --- | --- | --- | --- | --- | --- |
| 25 | 20 | 0.05 | 10.90 | 0.33 | 0.00 | 0.03 | 9.20 | 1.62 | 22.12 ± 5.7 |
| 26 | 13 | 2.08 | 33.81 | 2.85 | 0.65 | 0.12 | 63.69 | 7.65 | 110.85 ± 15.9 |
| 27 | 2 | 5.25 | 26.00 | 5.75 | 7.50 | 2.50 | 97.50 | 10.50 | 155.00 ± 42.5 |
| 28 | 17 | 14.24 | 52.24 | 14.76 | 12.74 | 6.71 | 125.24 | 12.76 | 238.68 ± 20.4 |
| 29 | 25 | 22.36 | 64.96 | 23.52 | 26.24 | 10.30 | 153.64 | 16.24 | 317.26 ± 11.8 |
| 31 | 1 | 21.00 | 91.00 | 29.00 | 45.00 | 26.50 | 141.00 | 17.00 | 370.50 |
| 32 | 1 | 28.00 | 89.00 | 35.00 | 56.50 | 34.50 | 179.00 | 12.00 | 434.00 |

\* one side counted, values presented assume symmetry

